# Using transfer learning and dimensionality reduction techniques to improve generalisability of machine-learning predictions of mosquito ages from mid-infrared spectra

**DOI:** 10.1101/2022.07.26.501594

**Authors:** Emmanuel P. Mwanga, Doreen J. Siria, Joshua Mitton, Issa H. Mshani, Mario Gonzalez Jimenez, Prashanth Selvaraj, Klaas Wynne, Francesco Baldini, Fredros O. Okumu, Simon A. Babayan

## Abstract

Accurate prediction of mosquito population age structures can improve the evaluation of mosquito-targeted interventions since old mosquitoes are more likely to transmit malaria than young ones. Mid-infrared spectroscopy (MIRS) reveals age-associated variation in the biochemical composition of the mosquito cuticle, which can then be used to train machine learning (ML) models to predict mosquito ages. However, these MIRS-ML models are not always generalisable across different mosquito populations. Here, we investigated whether dimensionality reduction applied to the MIRS input data and transfer learning could improve the generalisability of MIRS-ML predictions for mosquito ages. We reared adults of the malaria vector, *Anopheles arabiensis*, in two insectaries (Ifakara, Tanzania and Glasgow, UK). The heads and thoraces of female mosquitoes of two age classes (1-9 day-olds and 10-17 day-olds) were scanned using an attenuated total reflection-Fourier transform infrared (ATR-FTIR) spectrometer (4000 cm^-1^ to 400 cm^-1^). The dimensionality of the spectra data was reduced using unsupervised principal component analysis (PCA) or t-distributed stochastic neighbour embedding (t-SNE), and then the spectra were used to train deep learning (DL) and standard machine learning (ML) classifiers. Transfer learning was also evaluated for improving the computational cost of the models when predicting mosquito age classes from new populations. Model accuracies for predicting the age of test mosquitoes from the same insectary as the training samples reached 99% for DL and 92% for ML, but did not generalise to a different insectary, achieving only 46% and 48% for ML for DL, respectively. Dimensionality reduction did not improve the model generalisability between locations but reduced computational time up to 5-fold. However, transfer learning by updating pre-trained models with 2% of mosquitoes from the alternate location brought both DL and standard ML model performance to ~98% accuracy for predicting mosquito age classes in the alternative insectary. Combining dimensionality reduction and transfer learning can reduce computational costs and improve the transferability of both deep learning and standard machine learning models for predicting the age of mosquitoes. Future studies could investigate the optimal quantities and diversity of training data necessary for transfer learning, and implications for broader generalisability to unseen datasets.

## Background

Malaria currently kills approximately one child every minute [1]. In 2020, there were 241 million cases and 627,000 deaths, nearly all in Sub-Saharan Africa [1]. Currently, the most widespread and cost-effective method of malaria prevention is based on controlling the mosquitoes that transmit the disease. Since 2000, insecticide-treated nets (ITNs) and indoor residual spraying (IRS) have so far contributed nearly 80% of all global malaria decline [2]. However, the direct impact of individual control programs on the mosquito populations and on malaria transmission at the sites of intervention remains difficult to measure. To guide further efforts against the disease, evaluating the performance of these and other vector control interventions is crucial for measuring their impact in different settings. The World Health Organization (WHO) now recommends that surveillance be integrated as a core component of malaria control programs [3].

This necessitates scalable, simple-to-implement and low-cost methods for quantifying key biological attributes of mosquitoes, such as age, infection status, and blood meal preferences, which are essential for understanding pathogen transmission dynamics. The age and survivorship of key *Anopheles* vectors are especially important in determining the likelihood that the mosquitoes will live long enough to allow complete parasite development (the extrinsic incubation period), and subsequent transmission to humans [4]. The assessments are essential for monitoring the impacts of interventions such as ITNs and IRS, which primarily kill adult mosquitoes in the field [5].

The current “gold standard” for estimating the age of malaria mosquitoes is to dissect their ovaries to estimate how many times they have laid eggs [5,6]. Despite their low technical demands, such procedures are time-consuming and labour-intensive. Age-grading dissections can also be imprecise because of gonotrophic discordance, which is common in Afrotropical malaria vectors [7], or of their reliance on the availability of host blood meals, which determines when and how frequently a mosquito blood-feeds.

We and others have demonstrated that spectroscopic analysis of mosquitoes using near infrared (12500 – 4000 cm^-1^) or mid-infrared (MIR) (4000 – 400 cm^-1^) frequencies can identify key biochemical signals that vary with age [8,9]. These methods, when combined with specific machine learning (ML) techniques, allow for rapid estimation of mosquito ages [9,10].

Despite early successes, these infrared-based applications have limitations such as their portability to mosquitoes from different locations or laboratories [10] and the substantial computational requirements for retraining such models. Indeed, the inherent variability of mosquitoes from different environmental and genetic backgrounds may limit the generalisability of models trained on infrared spectra. The models could also be misled by signals in MIR that are associated with confounding factors introduced during sampling (e.g., atmospheric contamination with water vapour, temperature variations and high humidity in the laboratory), thus learning features that are not strictly related to the biochemical trait being investigated. As a consequence, models currently must be regularly retrained using new data from target mosquito populations.

To increase the generalisability of ML models for a given training dataset, a variety of spectral smoothing and regularisation techniques have been tested, such as penalised regression [11]. These methods are known to be computationally efficient and to improve generalisability [11]. Deep learning (DL) techniques such as convolutional neural networks (CNN) have recently been used on large spectra data [10], improving generalisability through transfer learning (i.e., updating a pre-trained model with a small amount of new data from a different target population). However, when trained on large datasets, such techniques remain computationally expensive and may necessitate repeated sampling of hundreds of mosquitoes from different populations and environments to allow successful generalisability. Alternatively, since standard ML models are less complex than DL, computational time can be kept to a minimum. DL methods are versatile extensions of machine learning that are ideal for complex or large datasets [12]. But are prone to overfitting (predicting the training dataset well but failing on previously unseen or new data).

However, unsupervised learning algorithms, which find patterns independent of pre-defined target labels, can aggregate, cluster or eliminate features while retaining dominant statistical information before training. The resulting dimensionality reduction may improve generalisability as well as reducing computational costs of training models. Examples include principal component analysis (PCA) [15–17], which projects a large number of variables into distinct categories that summarise data into a small number of independent principal components, and t-distributed Stochastic Embedding (t-SNE) [18], which clusters datapoints based distances between all their input dimensions.

This study assessed whether the generalisability and computational costs of MIRS-based models for predicting the age classes of female *An. arabiensis* mosquitoes reared in two different insectaries in two locations could be improved by combining dimensionality reduction and transfer learning methods.

## Methods

### Collection of mosquito spectra data

We analysed mid-infrared spectra from two strains of *An. arabiensis* mosquitoes obtained from two different insectaries, one from University of Glasgow, UK and another from Ifakara Health Institute, Tanzania. The same data had previously been used to demonstrate the capabilities of mid-infrared spectroscopy and CNN for distinguishing between species and determining mosquito age [10]. The insectary conditions under which the mosquitoes were reared (temperature 27 ± 1.0 °C, and relative humidity 80 ± 5%) have been described elsewhere [19].

Mosquitoes were collected from day 1 to day 17 after pupal emergence at both laboratories and divided in two age classes (1-9 day-olds and 10-17 day-olds). Silica gel was used to dry the mosquitoes. For each chronological age in each laboratory, ~120 samples were measured by MIRS on each day. The heads and thoraces of the mosquitoes were then scanned with an attenuated total reflectance Fourier-Transform Infrared (FTIR) ALPHA II and Bruker Vertex 70 spectrometers both equipped with a diamond ATR accessory (BRUKER-OPTIC GmbH, Ettlingen, Germany). The scanning was performed in the mid-infrared spectral range (4000 – 400 cm^-1^) at a resolution of 5 cm^-1^, with each sample being scanned 16 times to obtain averaged spectra as previously described [9,20].

### Data pre-processing

The spectral data were cleaned to eliminate bands of low intensity or significant atmospheric intrusion using the custom algorithm [21]. The final datasets from Ifakara and Glasgow contained 1720 and 1635 mosquito spectra, respectively. In these two datasets, the chronological age of *An. arabiensis* was categorised as 1 - 9 days old (i.e. young mosquitoes representative of those typically unable to transmit malaria) and 10–17 days old (i.e. older mosquitoes representative of those potentially able to transmit malaria) [22].

To improve the accuracy and speed of convergence of subsequent algorithms, data were standardised by centring around the mean and scaling to unit variance [23].

### Dimensionality reduction

Principal component analysis (PCA) and t-distributed stochastic neighbour embedding (t-SNE) were used separately to reduce the dimensionality of the data [15–18]. Both PCA and t-SNE were implemented using the scikit-learn library [23].

Separately, t-SNE was used to convert high-dimensional Euclidean distances between spectral points into joint probabilities representing similarities. To cluster the data into three features, the embedded space was set to 3, because the Barnes-hut algorithm in t-SNE is limited to only 4 components. Perplexity was set to 30 as the number of nearest neighbours, which means that for each point, the algorithm took the 30 closest points and preserved the distances between them. For smaller datasets perplexity values ranging from 5 and 50 are thought to be optimal for avoiding local variations and merged clusters caused by small or large perplexity values [18]. The learning rate for t-SNE is generally in the range of 10 - 1000 [23], thus it was set to 200 scalar.

### Machine learning training

#### Deep learning

DL models were trained and used to classify the *An. arabiensis* mosquitoes into the two age classes (1-9 or 10-17 day-olds). The intensities of *An. arabiensis* mid-infrared spectra (matrix of features) were used as input data, while the model outputs were the mosquito age classes.

Three different deep learning models were trained; 1) Convolutional neural network (CNN) model without dimensionality reduction, 2) Multi-Layer Perceptron (MLP) with PCA as dimensionality reduction, and 3) MLP with t-SNE as dimensionality reduction. For all models, a SoftMax layer was added to transform the non-normalized outputs of *K*-units in a fully connected layer into a probability distribution of belonging to either one of two age classes (1–9 or 10–17 days). Moreover, to compute the gradient of the networks, stochastic gradient boosting was used as an optimisation algorithm [25], and categorical cross-entropy loss was used for the classifier’s metric.

To begin, we trained a one-dimensional CNN model with four convolutional layers and one fully connected layer when the dimensionality of the data was not reduced (Figure 1A), and therefore consisting of 1666 training features from the data. The one-dimensional CNN was used because it is effective at deriving features from fixed-lengths (i.e. the wavelengths of the mid-infrared spectra), and it has been previously been used efficiently with spectral data [19]. To extract features from spectral signals, the deep learning architecture used convolutional, max-pooled and fully connected layers. The convolutional operation was carried out with kernel sizes (window) of 8, 4, and 6, and a kernel window shift size (stride) of either 1 or 2. For each kernel size, 16 filters were used to detect and derive features from the input data. Furthermore, given the size of the training data, the fully connected layer consisted of 50 neurons to reduce the model’s complexity.

**Figure 1:**
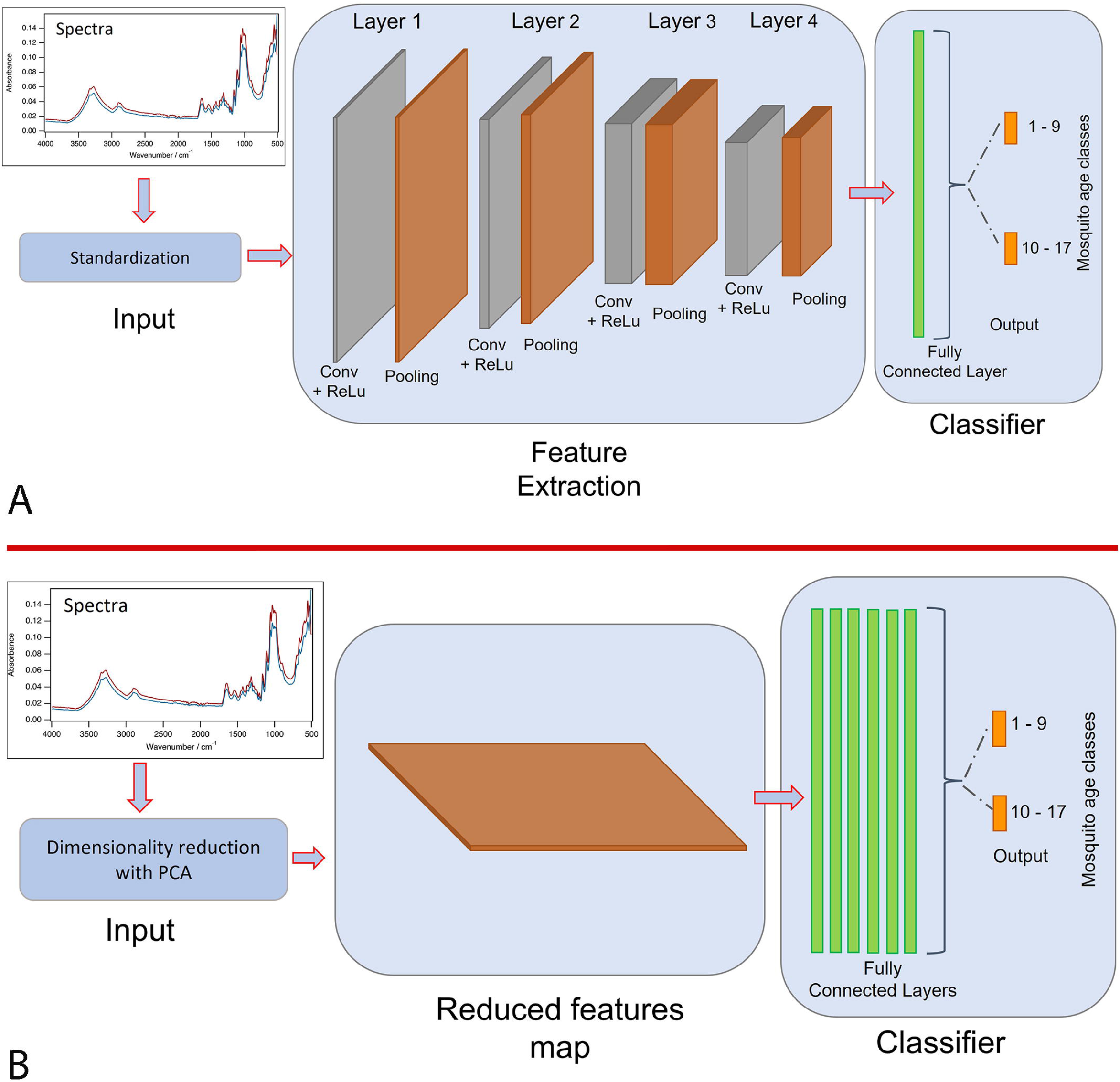
A schematic representation of a deep learning models that uses mosquito spectra as input to predict mosquito age classes. **A)** CNN - no dimensionality reduction is applied: standardised spectral features are fed as input through four different convolutional layers, followed by one fully connected layer, with the predicted age classes shown as the output layer. **B)** MLP - dimensionality reduction is used: spectral features that have been reduced in dimension using PCA or t-SNE are fed as input through 6 fully connected layers, with the predicted age classes shown as the output layer.

Moreover, batch normalisation layers were added to both models to improve model stability by keeping mean activation close to 0 and activation standard deviation close to 1. To reduce the likelihood of overfitting, dropout was used during model training to randomly and temporarily remove units from the network at a rate of 0.5 per step. Furthermore, after 50 rounds, early stopping was used to halt training when a validation loss stopped improving.

#### Dimensionality reduction

We trained two additional deep learning models, in this case Multi-Layer Perceptron (MLP), with PCA or t-SNE transformed input data (Figure 1B). The models were trained with only fully connected layers (n = 6) containing 500 neurons each, given the limited number of training features to ensure performance and stability. To control for overfitting, the procedure was similar to that of the CNN above, except that early stopping was used to halt training when a validation loss stopped improving after 500 rounds.

#### Transfer learning

The Ifakara dataset was used to pre-train the ML model. The Ifakara dataset was divided into training and test sets, and estimator performance was assessed using *K*-fold cross-validation (*k* = 5) [26], (Figure 3). We therefore determined what percentage of the new spectra data from the alternate location was required for ML models to sufficiently learn the variability in the other insectary. To put transfer learning options to the test, either 82 or 33 spectra were randomly selected from the 1635 of the Glasgow data, accounting for 5% and 2% of the dataset, respectively. The learning process in this case relied on a pre-trained model (trained with Ifakara data), avoiding the need to start training from scratch (Figure 2). The ML models pre-trained with Ifakara dataset were fine-tuned using 2% or 5% subsets of the Glasgow dataset. The output was compared to that of a model trained solely with Ifakara data (i.e., no transfer learning).

**Figure 2:**
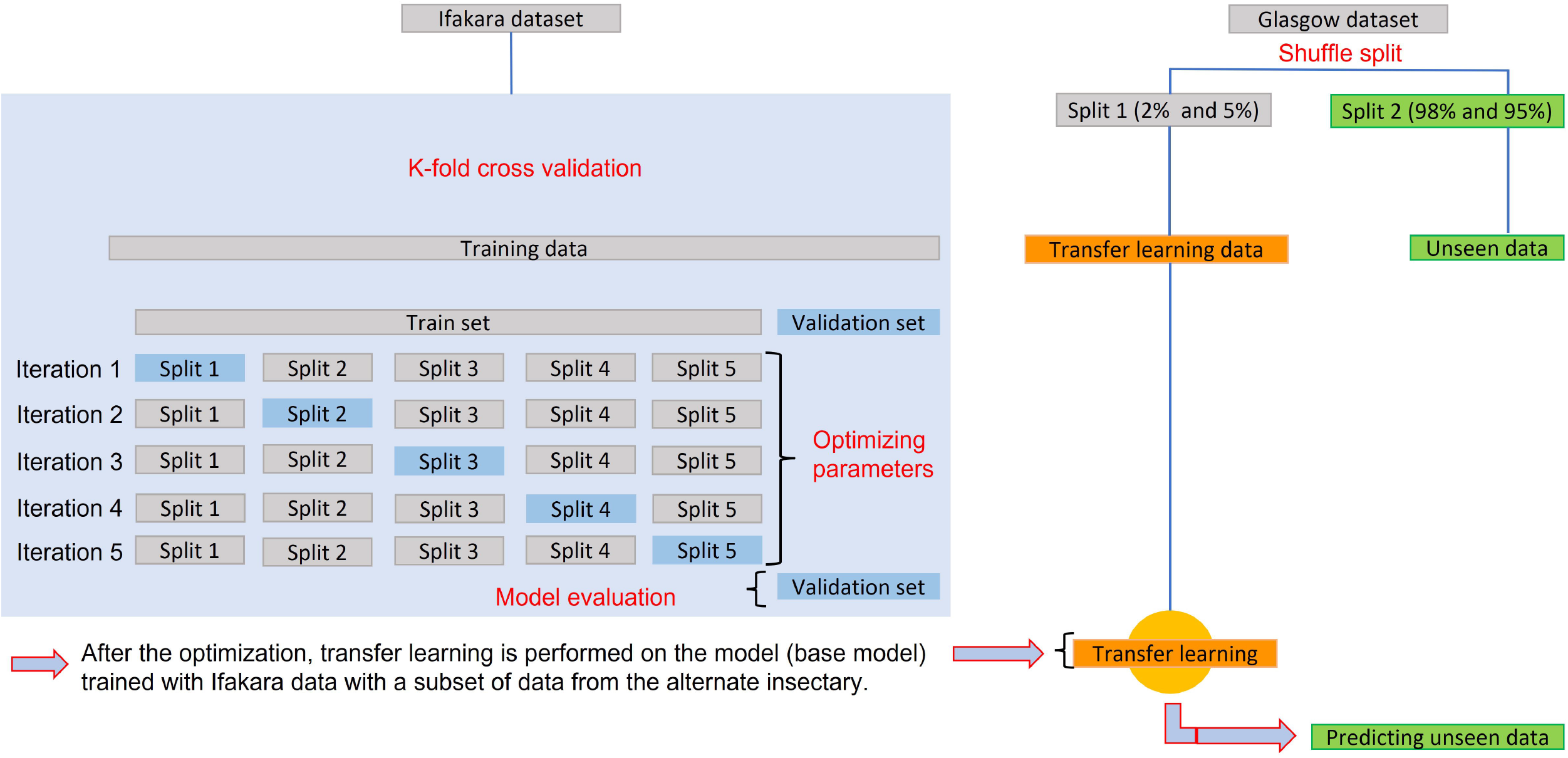
Schematic illustrating the process of data splitting, model training, cross-validation, and transfer learning.

**Figure 3:**
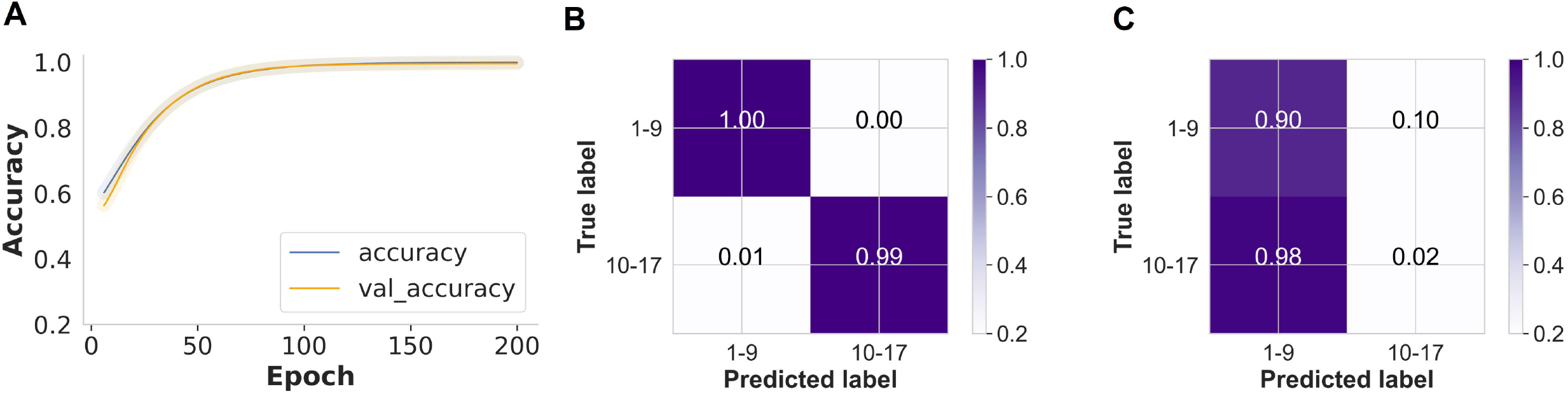
CNN generalisation and prediction of mosquito age using data from a single insectary (Ifakara) with no dimensionality reduction. **A**) Training and validation classification accuracy for mosquito age classes improved from ~60% to 95% as training iterations increased (200 epochs). **B)** A normalised confusion matrix displaying the proportions of correct mosquito age class predictions achieved on the held-out Ifakara data (test set) during model training. **C)** Proportions of correct mosquito age class predictions based on unseen data from the alternate insectary (Glasgow).

Precision, recall, and F1-scores were calculated from predicted values for each age class to demonstrate the validity of the final models in predicting the unseen Glasgow data. Keras and TensorFlow version 2.0 were used for deep learning process [27,28].

#### Standard machine learning

We also compared the prediction accuracy of CNN to that of a standard machine learning model trained on spectra data transformed by PCA or t-SNE. Different algorithms were compared, including K-Nearest Neighbour, logistic regression, support vector machine classifier, random forest classifier, and a gradient boosting (XGBoost) classifier. The model with the highest accuracy score for predicting mosquito age classes was optimised further by tuning its hyper-parameters with randomised search cross-validation [23]. The cross-validation evaluation used to assess estimator performance in this case was the same as that used in deep learning. The fine-tuned model was used to predict mosquito age classes in previously unseen Glasgow dataset.

Python version 3.8 was used for both the deep learning and standard machine learning training. All computations were done on a computer equipped with 32 Gigabits (gb) of random-access memory (RAM) and an octa-core central processing unit.

## Results

### DL mosquito age classification with and without dimensionality reduction, did not generalise between the two locations

In the initial analysis, only spectra from the Ifakara insectary were used to train the CNN. During model training, the CNN classifier achieved 99% training accuracy without any dimensionality reduction (Figure3A). When given new held-out data from the same Ifakara insectary (test set), the model predicted mosquitoes aged 1 – 9 days with 100% accuracy and those aged 11 – 17 days with 99% accuracy (Figure 3B). However, when the same model was used to predict age classes for Glasgow insectary samples, the overall accuracy was 46%, and therefore indistinguishable from any random classifications (Figure 3C).

In addition, a CNN classifier required 200 epochs for training, with a running time of 7.2 – 7.8 seconds per epoch when no dimensionality reduction on the input data was used (Table 1).

**Table 1:**
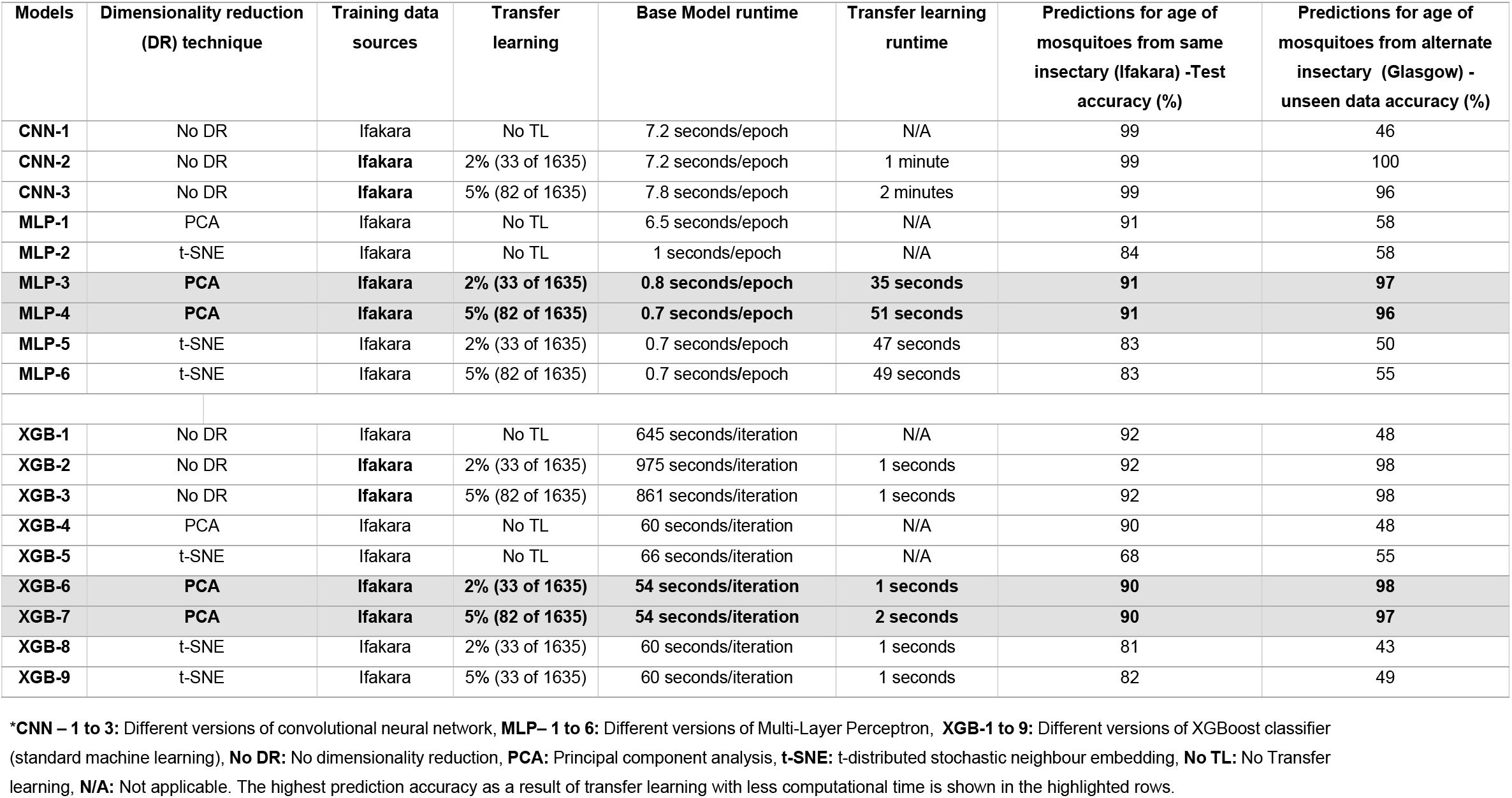
The performance of deep learning and standard machine learning models for predicting mosquito age classes from the same or alternate insectaries, with and without dimensionality reduction (DR) and transfer learning.

When PCA was used to reduce the dimensionality of the data, the MLP model trained with only Ifakara spectra predicted the held-out data from the same insectary (Ifakara) with an overall accuracy of 91%, but could attain only 58% accuracy for predicting age classes of Glasgow mosquitoes (Table 1). Similarly, when t-SNE was used as the dimensionality reduction technique, the model predicted the held-out Ifakara data (test set) with an accuracy of 85%, but failed to accurately predict age classes of Glasgow data (Table 1).

Furthermore, when PCA or t-SNE were used to transform the input data, a MLP classifier needed 5000 epochs to train, with a running time of 0.7 - 0.8 seconds per epoch (Table 1)

### Transfer learning improves DL accuracy and generalisability

To improve generalisability (i.e., the ability of the models to predict the age classes of samples from other sources), we tuned the pre-trained CNN models with 2% or 5% of the spectra from Glasgow (i.e., 2% or 5% target population samples for transfer learning), and used the updated model to predict the unseen Glasgow dataset. When no dimensionality reduction was used, the pre-trained model predicted the held-out test (Ifakara dataset) with 99% accuracy and transferred well to the Glasgow dataset when 2% and 5% target population samples were used for transfer learning, achieving 100% and 96% accuracies, respectively (Table 1).

However, when PCA or t-SNE were used to reduce the dimensionality of the data, the MLP classifier was trained with only fully connected layers in this case to allow the model to learn the combination of features with the network’s learnable weights. Using PCA, the pre-trained model predicted the held-out test (Ifakara dataset) with 91% accuracy, but when 2% transfer learning was applied, the model transferred well to the Glasgow dataset, achieving 97% accuracy when predicting the mosquito age classes, and 96% accuracy with 5% target population samples (Table 1, Figure 5A-C).

**Figure 4:**
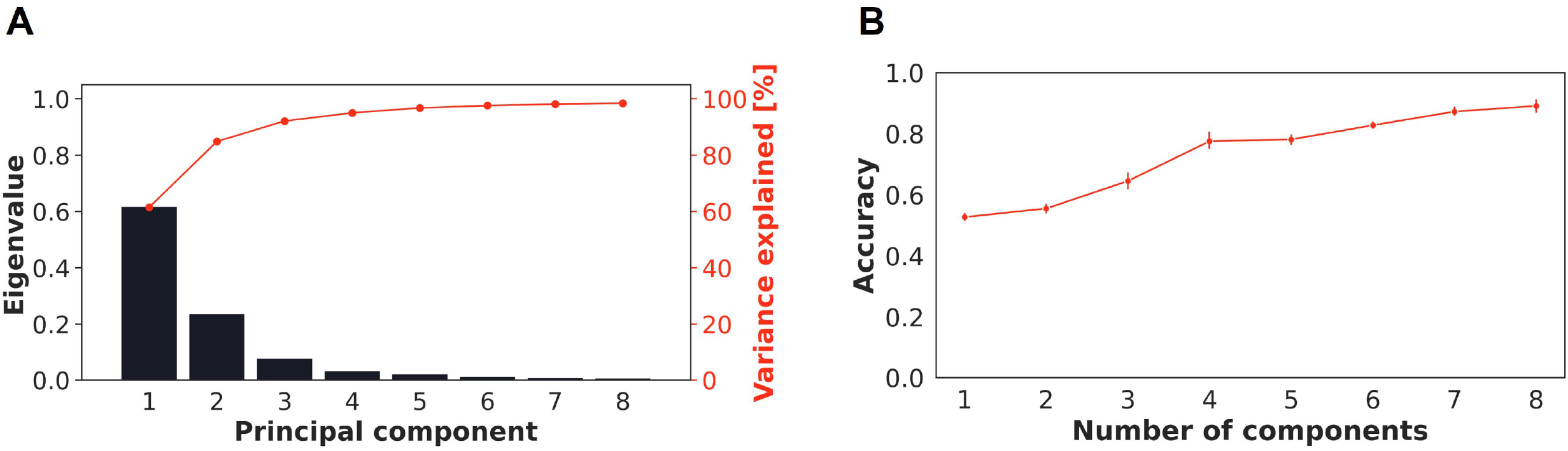
**A)** cumulative explained variance and eigenvalues as the function of principal components. **B)** Number of principal components included in the XGB classifier (i.e. from 1:8 PCs).

**Figure 5:**
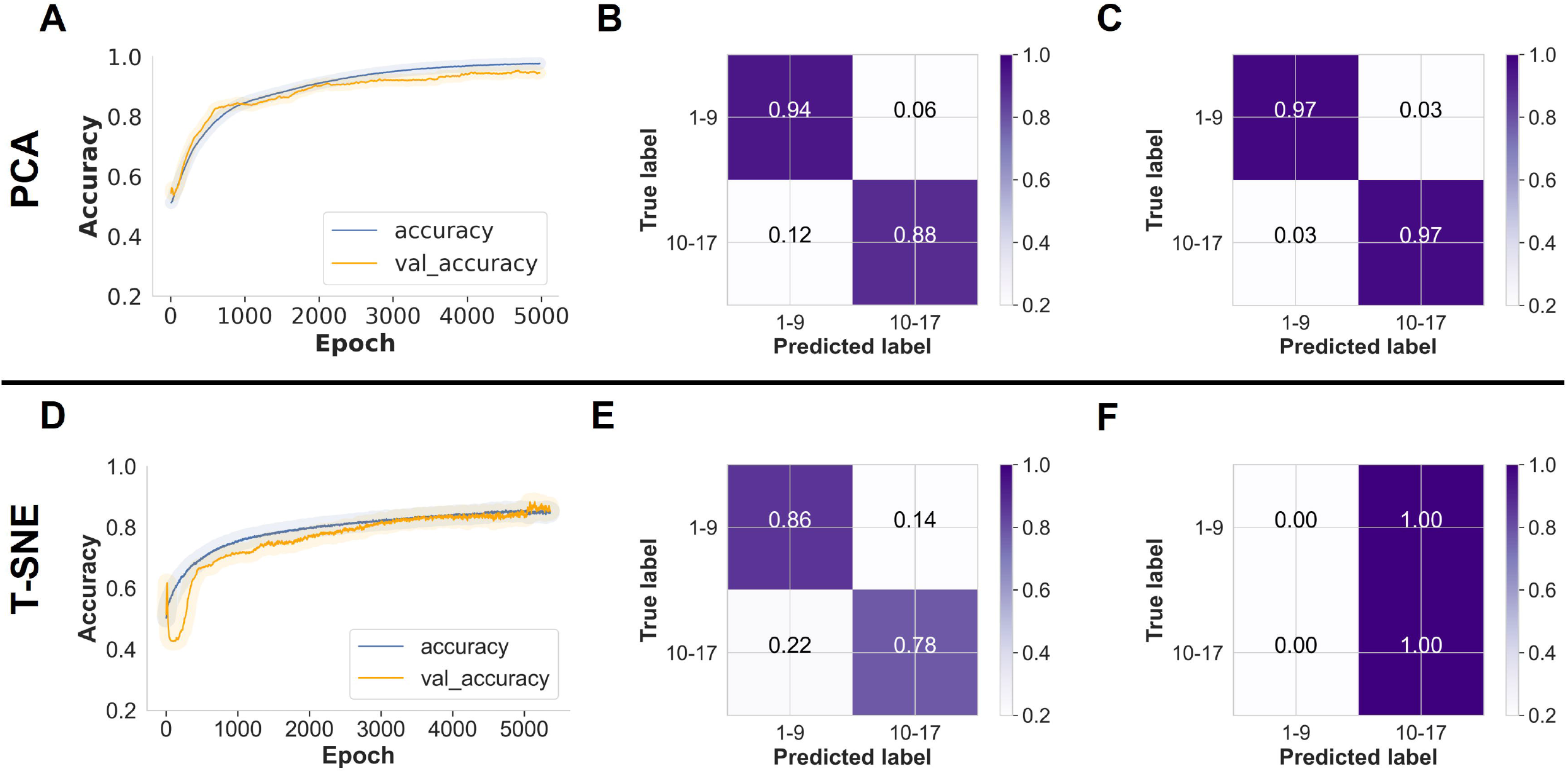
MLP trained on PCA-transformed Ifakara dataset plus 2% new target population samples: **A**) As training time increased (5000 epochs), training and validation classification accuracy for mosquito age classes increased from 50% to 91%, **B)** A normalised confusion matrix displaying the proportions of correct mosquito age class predictions achieved on the held-out Ifakara test set during model training, **C)** Proportions of correct mosquito age class predictions achieved on unseen Glasgow dataset. **MLP trained on t-SNE-transformed Ifakara dataset plus 2% new target population samples: D)** As training time increased (5000 epochs), training and validation classification accuracy for mosquito age classes increased from 60% to 83%, **E)** A normalised confusion matrix displaying the proportions of correct mosquito age class predictions achieved on the held-out Ifakara test set during model training, **F)** Proportions of correct mosquito age class predictions achieved on unseen Glasgow dataset.

PCA was used to project the data into lower dimensional space using singular value decomposition [15,24], with the goal of achieving the best summary using optimal number of principal components (PCs) with up to 98% of variance explained (Figure 4A). Further, when the impact of PCs on accuracy was assessed, a greater prediction accuracy was found, leading to the selection of 8 PCs. (Figure 4B).

When using t-SNE, the pre-trained predicted the age classes in the held-out data (test set) with 83% accuracy but failed to achieve generalisability for the Glasgow data when either 2% or 5% transfer learning was applied, achieving only 50% and 55% accuracy, respectively (Table 1, Figure 5D-F).

Transfer learning also reduced training time while improving the performance of both DL and standard machine learning models in predicting samples from the target population. Transfer learning took less than two minutes for both models to produce the desired results (Table 1).

### Comparison between deep learning and standard machine learning models in achieving generalisability

The XGBoost classifier, when trained with Ifakara data only, failed to predict age classes of mosquitoes from the Glasgow insectary, with or without dimensionality reduction (Table 1). However, when the classifier was updated with 2% target population samples, the model correctly classified individual mosquito age classes with 98% for both 1–9 days old and 10– 17 days old mosquitoes (Figure 6). Similar results were observed even when PCA was used as a dimensionality reduction (Figure 6). Increasing the samples for transfer learning to 5% of the training set had no effect on the accuracies (Table 1). However, when t-SNE was used for dimensionality reduction, transfer learning with either 2% or 5% Glasgow samples did not improve the generalisability of the XGBoost classifier (Table 1).

**Figure 6:**
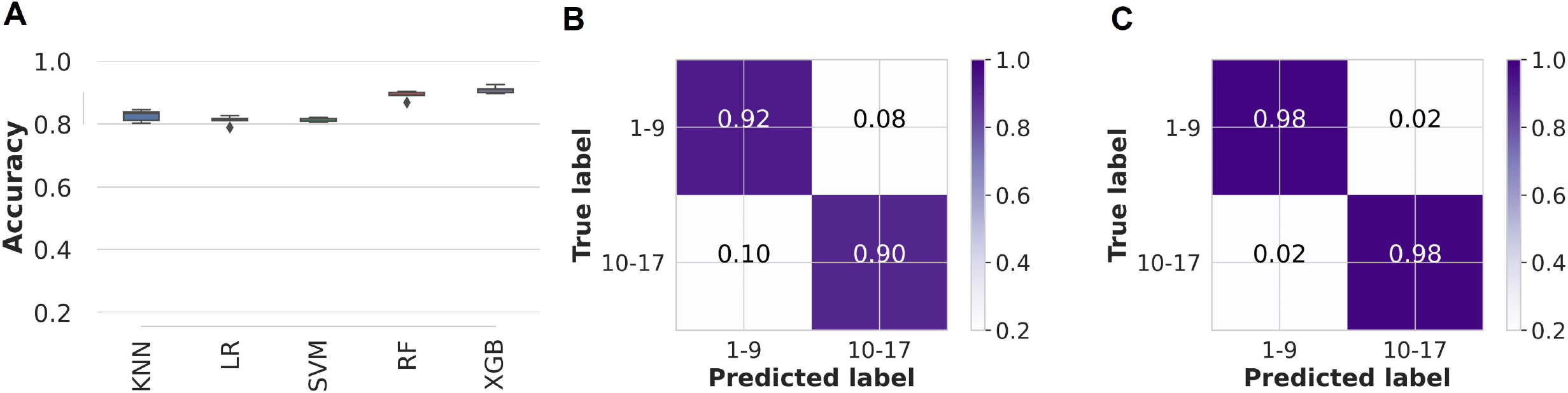
Standard machine learning models’ predictive accuracies and generalisability when trained with PCA-transformed Ifakara data plus 2% new target population. **A)** Comparison of standard machine learning models for mosquito age classification; KNN: K-nearest neighbours, LR: Logistic regression, SVM: Support vector machine classifier, RF: Random forest classifier, and XGB: XGBoost. **B)** A normalised confusion matrix displaying the proportions of correct mosquito age class predictions achieved on the held-out data (test set) during model training. **C)** proportions of correct mosquito age class predictions achieved on unseen Glasgow dataset.

Table 2 shows how the performance of deep learning and standard machine learning was evaluated using other metrics such as precision, recall, and f1-scores. When it comes to mosquito age classification, the XGBoost classifier matches the deep learning model in both specificity (precision) and sensitivity (recall).

**Table 2:** Precision, recall, and f1-score of the best deep learning model for classifying mosquito age classes from alternate sources compared to the best standard machine learning algorithm (i.e. XGBoost classifier).

| Model name | Age class (Days) | Precision | Recall | f-1 score | No. of samples per age class |
| --- | --- | --- | --- | --- | --- |
| **MLP-3** | 1-9 | 0.98 | 0.97 | 0.98 | 895 |
|  | 10-17 | 0.97 | 0.97 | 0.97 | 707 |
| **XGB-6** | 1-9 | 0.98 | 0.99 | 0.98 | 895 |
|  | 10-17 | 0.98 | 0.98 | 0.98 | 707 |

\* **MLP-3**: Multi-Layer Perceptron trained with PCA as a dimensionality reduction technique and 2% transfer learning, **XGB-6**: XGBoost classifier trained with PCA as a dimensionality reduction technique and 2% target population samples used for transfer learning.

Further to that, standard machine learning models were trained with 10 iterations, and still the computing runtimes were generally shorter than those for CNN models when PCA and t-SNE were used to transform the input data, in some cases by up to 5 times (Table 1).

## Discussion

This study demonstrates that transfer learning approaches can substantially improve the generalisability of both deep learning and standard machine learning in predicting the age class of mosquitoes reared in two different insectaries. We evaluated 1635 mosquito spectra from Glasgow-reared mosquitoes and show that using transfer learning and dimensionality reduction techniques could improve machine learning models to predict mosquito age classes from alternate insectaries. Furthermore, reducing the dimensionality of the spectral data reduced computational costs (i.e. computing time) when training the machine learning models.

The current study adds to the growing evidence of the utility of infrared spectroscopy and machine learning in estimating mosquito age and survival [8,29–31]. In the past, most applications of infrared spectroscopy in estimating mosquito vector survival relied on nearinfrared frequencies (12,500 cm^-1^ to 4000 cm^-1^). A recent study used mid-infrared spectra (from 4000 cm^-1^ to 400 cm^-1^ frequencies) and standard machine learning to distinguish mosquito species with up to 82% accuracy, but found lower age prediction accuracy in several alternate settings [9]. González *et al*., suggested that machine learning underprediction may be explained by the small training dataset and ecological variability between the training and validation sets [9].

In our study, despite categorising mosquito chorological age into two classes (young: 1-9-day olds and old: 10–17-day olds), deep learning and standard machine learning approaches both remained unable to generalise, even after reducing the dimensionality of the spectra data. This result is consistent with Siria *et al*. [10], where CNN underperformed as a result of the difference in data distribution between the training and evaluation data driven by non-genetic factors such as ecological variation. When near-infrared spectroscopy was used to predict the age of *Anopheles* mosquitoes reared from wild populations, a similar limitation was reported [8,29].

Nonetheless, Siria *et al*. [10] also observed that using transfer learning to correct the difference data distribution between training and evaluation data improved deep learning generalisation, achieving 94% accuracy in predicting both species and mosquito age classes. Furthermore, in the latter study, the performance of the classifier was improved by incorporating a subset (*n* = 1200 ~ 1300 spectra) of the evaluation data into the training data.

The present study shows performing transfer learning using 2% of the spectra from the target domain (33 of 1635) as well as dimensionality reduction resulted in the improved generalisability of both deep learning and standard machine learning models achieving overall accuracy of ~98%. In this case, we expected that all models to which transfer learning was applied would outperform the baseline models. However, as the proportion of data from the target domain in the training increased, the performance slightly dropped for the deep learning. The reason for the deterioration in performance after turning the pre-trained/base model with 5% transfer learning could be that the model overfitted random noise during training, which negatively impacted the performance of these models on unseen data. Other studies have proposed alternative transfer learning approaches, such as adaptative regularisation to address cross-domains (i.e. source domain and target domain) learning problems [32], transferring knowledge gained in the source domain during training to the target domain [33], and integrating dimensionality reduction to transform features of the source to ensure data distribution in different domains is minimised [34].

Furthermore, dimensionality reduction was used in conjunction with transfer learning to reduce noise, redundant features, and computational time. Based on our findings, dimensionality reduction alone cannot achieve generalisability of machine learning models. The PCA improved model stability because the eigenvectors of the correlation matrix in PCA provide new axes of variation to project new data. The model with t-SNE as a dimensionality reduction technique failed to achieve generalisability on the new data, the reason for poor performance could be t-SNE is a probabilistic technique with a non-convex cost function [18], causing the output to differ from multiple runs.

Furthermore, incorporating dimensionality reduction substantially reduces model training time and thus, computational requirements. When compared to models trained without dimensionality reduction, the computing runtimes for models trained with dimensionality reduction were less than five-fold. Moreover, transfer learning in general was fast, tuning the pre-trained models in under two minutes on our machine (standard laptop). This makes the technique applicable and reproducible even to users with low computing power and capacity providing they have access to pre-trained models.

This study only included *An. arabiensis* reared in the laboratory from two insectaries. Future research should put the techniques to the test with samples from more laboratories, field settings, and mosquito species, as these factors can affect the model’s predictive capacity. The optimal ratio of transfer learning data required to achieve best generalisability in predicting mosquito age class has yet to be determined, so future studies could investigate this gap. Furthermore, because dimensionality reduction reduced the computational requirements in this study, we suggest that clustering spectra with algorithms such as PCA can be a beneficial strategy for models trained on MIRS.

## Conclusion

Dimensionality reduction does not improve model generalisability in predicting mosquito age classes using data from alternate insectary. However, using transfer learning and dimensionality reduction with PCA, the generalisability of the deep learning models in predicting mosquito age classes improved from 56% to >95%. Therefore, this study indicates that these techniques could be scaled up and evaluated further to determine the age of mosquitoes from different populations, transfer learning is currently necessary. Moreover, when dimensionality reduction and transfer learning are used, standard machine learning (e.g., the XGBoost classifier) can reduce computational time while also matching the performance of deep learning. The development of these models could reduce the amount of work and time required for entomologists to dissect large numbers of mosquitoes. These approaches could be used to improve model-based surveillance programmes, such as assessing the impact of malaria vector control tools, by monitoring the age structures of local vector populations.

## Abbreviations

CNN: Convolutional neural network
ITNs: Insecticide treated nets
PCA: Principal component analysis
t-SNE: t-distributed stochastic neighbour embedding

## Acknowledgement

The authors express their gratitude to for the research and administrative support teams at Ifakara Health Institute and the University of Glasgow.

## Contributions

EPM, SAM, DJS, FB, MGJ and FOO designed the study. DJS supported in data collection semi-field experiments. EPM performed data analysis. JM provided technical support to EPM during data analysis. EPM wrote and revised the manuscript. EPM, SAB, IHM, PS, FOO, and FB reviewed and revised the manuscript. All authors read and approved the final manuscript.

## Ethical clearance

At IHI, ethical approval for the study was obtained from the IHI Institutional Review Board (Ref. IHI/IRB/EXT/No: 005-2018), and from the National Institutes of Medical Research (NIMR), Ref: NIMR/HQ/R.8c/Vol.II/880. At the UofG, Ethical approval for the supply and use of human blood for mosquito feeding was obtained from the Scottish National Blood Transfusion Service committee for governance of blood and tissue samples for non-therapeutic use (submission Reference No 1815).

## Consent for publication

Permission to publish this work was also obtained from the National Institutes of Medical Research (NIMR) Ref. No: NIMR/HQ/P.12 VOL XXXIV/69.

## References

1. Geneva: World Health Organization. World Malaria report 2021. 2021.

2. Bhatt S, Weiss DJ, Cameron E, Bisanzio D, Mappin B, Dalrymple U, et al. The effect of malaria control on Plasmodium falciparum in Africa between 2000 and 2015. Nature. 2015;

3. WHO, World Health Organization, World Health Organization. Global Malaria Programme. Global technical strategy for malaria 2016-2030 [Internet]. World Heal. Organ. 2015. Available from: http://apps.who.int/iris/bitstream/10665/176712/1/9789241564991_eng.pdf?ua=1

4. Charlwood JD, Smith T, Billingsley PF, Takken W, Lyimo EOL, Meuwissen JHET. Survival and infection probabilities of anthropophagic anophelines from an area of high prevalence of Plasmodium falciparum in humans. Bull Entomol Res. 1997;87:445–53.

5. Detinova TS. Age-grouping methods in Diptera of medical importance with special reference to some vectors of malaria. Monogr Ser World Health Organ. Geneva: World Health Organization; 1962;

6. P. P V. The determination of the physiological age of female Anopheles by the number of gonotrophic cycles completed. Medskaya Parazit [Internet]. 1949 [cited 2022 Jul 9];18:352–5. Available from: https://cir.nii.ac.jp/crid/1572824499775933056.bib?lang=en

7. Rao V. On Gonotrophic Discordance among certain Indian Anopheles. Indian J Malariol. 1947;1:43–50.

8. Mayagaya VS, Michel K, Benedict MQ, Killeen GF, Wirtz RA, Ferguson HM, et al. Non-destructive determination of age and species of Anopheles gambiae sl using near-infrared spectroscopy. Am J Trop Med Hyg. 2009;81:622.

9. Gonzalez-Jimenez M, Babayan SA, Khazaeli P, Doyle M, Walton F, Reedy E, et al. Prediction of malaria mosquito species and population age structure using mid-infrared spectroscopy and supervised machine learning. Wellcome Open Res. 2019;4:76.

10. Siria DJ, Sanou R, Mitton J, Mwanga EP, Niang A, Sare I, et al. Rapid age-grading and species identification of natural mosquitoes for malaria surveillance. Nat Commun [Internet]. 2022;13:1501. Available from: https://doi.org/10.1038/s41467-022-28980-8

11. Esperança PM, Da DF, Lambert B, Dabiré RK, Churcher TS. Functional data analysis techniques to improve the generalizability of near-infrared spectral data for monitoring mosquito populations. bioRxiv [Internet]. 2020;2020.04.28.058495. Available from: http://biorxiv.org/content/early/2020/04/29/2020.04.28.058495.abstract

12. Géron A. Hands-On Machine Learning with Scikit-Learn and TensorFlow. First Edit. Tache N, editor. Boston: O’Reilly Media, Inc.; 2017.

13. Prechelt L. Early stopping - But when? Lect Notes Comput Sci (including Subser Lect Notes Artif Intell Lect Notes Bioinformatics). 2012;

14. Nitish S, Geoffrey H, Alex K, Ilya S, Ruslan S. Dropout: A Simple Way to Prevent Neural Networks from Overfitting. J Mach Learn Res. 2014;

15. Wold S, Esbensen K, Geladi P. Principal component analysis. Chemom Intell Lab Syst. 1987;

16. Lever J, Krzywinski M, Altman N. Principal component analysis. Nat Methods [Internet]. 2017;14:641–2. Available from: https://doi.org/10.1038/nmeth.4346

17. Schölkopf B, Smola A, Müller K-R. Kernel principal component analysis. Int Conf Artif neural networks. Berlin, Heidelberg: Springer; 1997. p. 583–8.

18. Van Der Maaten LJP, Hinton GE. Visualizing high-dimensional data using t-sne. J Mach Learn Res. 2008;

19. Siria DJ, Sanou R, Mitton J, Mwanga EP, Niang A, Sare I, et al. Rapid ageing and species identification of natural mosquitoes for malaria surveillance. bioRxiv [Internet]. 2020;2020.06.11.144253. Available from: http://biorxiv.org/content/early/2020/06/12/2020.06.11.144253.abstract

20. Mwanga EPP, Mapua SAA, Siria DJJ, Ngowo HSS, Nangacha F, Mgando J, et al. Using mid-infrared spectroscopy and supervised machine-learning to identify vertebrate blood meals in the malaria vector, Anopheles arabiensis. Malar J [Internet]. 2019;18:187. Available from: https://doi.org/10.1186/s12936-019-2822-y

21. Jiménez MG. A custom program that imports the IR spectra, cleans and screens them to eliminate the badly measured ones, and extracts the most interesting data from them! [Internet]. Wellcome Open Res. 2019. p. 4:76. Available from: https://github.com/SimonAB/Gonzalez-Jimenez_MIRS/blob/v1.0/Locomosquito.ipynb

22. Ohm JR, Baldini F, Barreaux P, Lefevre T, Lynch PA, Suh E, et al. Rethinking the extrinsic incubation period of malaria parasites. Parasit Vectors [Internet]. 2018;11:178. Available from: https://doi.org/10.1186/s13071-018-2761-4

23. Pedregosa F, Gramfort A, Michel V, Thirion B, Grisel O, Blondel M, et al. Scikit-learn: Machine Learning in Python. J Mach Learn Res. 2011;

24. Halko N, Martinsson PG, Tropp JA. Finding Structure with Randomness: Probabilistic Algorithms for Constructing Approximate Matrix Decompositions. SIAM Rev [Internet]. Society for Industrial and Applied Mathematics; 2011;53:217–88. Available from: https://doi.org/10.1137/090771806

25. Sutskever I, Martens J, Dahl G, Hinton G. On the importance of initialization and momentum in deep learning. 30th Int Conf Mach Learn ICML 2013. 2013.

26. Kohavi R. A Study of Cross-Validation and Bootstrap for Accuracy Estimation and Model Selection. Int Jt Conf Artif Intell. 1995;

27. Chollet F. Keras: The Python Deep Learning library. KerasIo. 2015;

28. Abadi M, Barham P, Chen J, Chen Z, Davis A, Dean J, et al. TensorFlow: A system for large-scale machine learning. Proc 12th USENIX Symp Oper Syst Des Implementation, OSDI 2016. 2016.

29. Lambert B, Sikulu-Lord MT, Mayagaya VS, Devine G, Dowell F, Churcher TS. Monitoring the Age of Mosquito Populations Using Near-Infrared Spectroscopy. Sci Rep. 2018;8.

30. Sikulu-Lord MT, Devine GJ, Hugo LE, Dowell FE. First report on the application of nearinfrared spectroscopy to predict the age of Aedes albopictus Skuse. Sci Rep. 2018;8.

31. Ntamatungiro AJ, Mayagaya VS, Rieben S, Moore SJ, Dowell FE, Maia MF. The influence of physiological status on age prediction of Anopheles arabiensis using near infrared spectroscopy. Parasites and Vectors [Internet]. 2013;6:298. Available from: http://parasitesandvectors.biomedcentral.com/articles/10.1186/1756-3305-6-298

32. Long M, Wang J, Ding G, Pan SJ, Yu PS. Adaptation Regularization: A General Framework for Transfer Learning. IEEE Trans Knowl Data Eng. 2014;26:1076–89.

33. Si S, Tao D, Geng B. Bregman Divergence-Based Regularization for Transfer Subspace Learning. IEEE Trans Knowl Data Eng. 2010;22:929–42.

34. Pan SJ, Tsang IW, Kwok JT, Yang Q. Domain Adaptation via Transfer Component Analysis. IEEE Trans Neural Networks. 2011;22:199–210.

